# Near infrared radiation-driven oxygenic photosynthesis contributes substantially to primary production in biofilms harboring chlorophyll *f*-containing cyanobacteria

**DOI:** 10.1101/2024.04.02.587864

**Authors:** Maria Mosshammer, Erik C. L. Trampe, Niels-Ulrik Frigaard, Michael Kühl

## Abstract

Cyanobacteria with far-red light photoacclimation (FaRLiP) can modify their photopigmentation by synthesizing red-shifted phycobiliproteins and chlorophylls, i.e., chlorophyll (Chl) *d* and *f*. This enables use of near-infrared radiation (NIR) for oxygenic photosynthesis in habitats depleted of visible light (VIS). Cyanobacteria with FaRLiP are widespread but their quantitative importance for primary production in natural habitats remains unknown. Previously we showed that intertidal beachrock formations can harbor endolithic populations of Chl *f*-containing cyanobacteria capable of using NIR for oxygenic photosynthesis (Kühl et al., 2020). Here we use a combination of gas exchange measurements and luminescence lifetime-based O_2_ imaging to quantify how endolithic cyanobacteria with far-red chlorophylls contribute to the primary production of an intertidal beachrock habitat when exposed to a natural gradient of visible and near-infrared radiation. While VIS-driven photosynthesis predominantly took place in the dense cyanobacterial surface biofilm of beachrock, NIR-driven photosynthesis was mainly confined to a subsurface layer in the beachrock containing endolithic cyanobacteria with Chl *f* and *d*. Yet such subsurface, NIR-driven photosynthesis provided a substantial O_2_ production reaching >20% of the gross photosynthesis rates under comparable photon irradiance of visible light. This points to a hitherto overlooked role of far-red light acclimated cyanobacteria for primary production in natural habitats characterized by steep attenuation of visible light and relative enrichment in near-infrared radiation.

## INTRODUCTION

Many cyanobacteria can remodel their photosynthetic apparatus via synthesis of far red-shifted phycobiliproteins and chlorophylls (Chl), i.e., Chl *d* and *f*, when grown under near-infrared radiation (NIR) (Gan et al. 2015a). Such far-red light photoacclimation (FaRLiP; cf. Gan et al. 2014) has been found across all 5 domains of cyanobacteria (Gan et al., 2015b) and is regulated at the transcriptional level via a 21-gene cluster (Gan et al. 2015b) that also contains genes for a red phytochrome-triggered control cascade of FaRLiP (Zhao et al. 2015). In contrast to cyanobacteria in the genus *Acaryochloris* that exhibit a constitutive expression of Chl *d* as their major photopigment (Myashita et al. 1995), cyanobacteria with FaRLiP respond dynamically to light quality (Gan et al. 2014) and can synthesize red-shifted phycobiliproteins, minor amounts of Chl *d*, as well as Chl *f*. FaRLiP involves the modification and inclusion of Chl *f* in both PSI and PSII reactions centers (Itos et al. 2015, Nürnberg et al. 2018; Gisriel, 2024).

Chl *f* was first discovered in the filamentous cyanobacterium *Halomicronema hongdechloris* (*H. hongdechloris*) isolated from a stromatolite (Chen et al., 2010, 2012). Studies of *H. hongdechloris* and several other cyanobacterial strains have provided detailed insight in to the genetic regulation, biochemical pathways, and photophysiological implications of FaRLiP and the associated structural modifications of the photosynthetic apparatus (e.g. Nürnberg et al. 2018; Gisriel et al., 2020; Solier et al., 2020; Mascoli et al. 2022) that facilitate use of near-infrared radiation (>700 nm) for oxygenic photosynthesis. In contrast, the ecological role of cyanobacteria with FaRLiP in natural habitats remains much less explored. Cyanobacteria with Chl *f* have been found in a variety of terrestrial (Behrendt et al., 2015, 2010; Zhang et al., 2019; Murray et al., 2022) and aquatic (e.g. Akutsu et al., 2011; Trampe and Kühl, 2016; Ohkubo and Miyashita, 2017) habitats. By use of a genetic marker, Antonaru et al. (2020) identified the presence of cyanobacteria capable of FaRLiP based on analyses of metagenomic datasets from environmental samples representative of many different terrestrial and aquatic habitats across different continents and climatic zones. This further supports the notion that cyanobacteria with FaRLiP are widespread and may play a hitherto unknown role in primary production, based on their capability to extend the spectral window for oxygenic photosynthesis into the near infrared region (700-780 nm) (Chen and Blankenship, 2011; Chen 2014). However, quantification of NIR-driven primary production in habitats harboring cyanobacteria with FaRLiP, and comparison to visible light-driven oxygenic photosynthesis is lacking.

Intertidal beachrock habitats are widespread sedimentary features of (sub)tropical coastlines (Vausdaukas et al., 2007) that can harbor endolithic cyanobacteria with Chl *f* growing under a dense surface biofilm of other cyanobacteria that strongly absorb visible light (Trampe and Kühl 2016). We recently showed a pronounced NIR-driven O_2_ production in the Chl *f*-containing layer of beachrock (Kühl et al. 2020), indicating a potential for cyanobacteria with FaRLiP to contribute substantially to primary production.

However, these first estimates were hampered by a lack of data on beachrock primary production, and the NIR-driven rates of photosynthesis were estimated from imaging O_2_ dynamics in small hot-spots of activity, while illuminating beachrock cross sections homogeneously with a single low irradiance, which may not be representative of the *in-situ* light levels and differs from the natural, vertical light gradients in beachrock.

In this research advance, we close this knowledge gap and provide a quantification of visible and near-infrared radiation driven photosynthesis in 3 different beachrock biofilm communities. Our study shows that NIR-driven gross photosynthesis by cyanobacteria with FaRLiP reaches high rates (up to >20% of areal photosynthesis rates driven by visible light at comparable photon irradiance) and thus can contribute significantly to the primary production of beachrock biofilms. The ecophysiological plasticity of cyanobacteria expressing FaRLiP can thus have important implications for the primary production of their natural habitats.

## RESULTS AND DISCUSSION

*Beachrock samples and pigmentation*. Previous studies across the beachrock platform on Heron Island revealed three characteristic biofilm communities predominated by cyanobacteria, as characterized by their surface coloration (black, brown and pink) and microbial community composition (Diez et al. 2007; Trampe and Kühl 2016; Kühl et al. 2020). The biofilms are characterized by a <1-2 mm thin, optically dense surface layer on top of a more loosely defined, greenish endolithic layer 1-4 mm below the surface of the beachrock (Fig. 1 – figure supplement 1). The green endolithic zone was predominated by blue-green cell aggregates of unicellular, *Chroococcidiopsis*-like cyanobacteria containing Chl *f,* confirming earlier findings (Trampe et al. 2016; Kühl et al. 2020).

Hyperspectral imaging of beachrock cross-sections showed spectral signatures of Chl *a* absorption (670-680 nm) in the surface layers, while the endolithic layers carried spectral signatures indicative of both Chl *a* and far-red absorbing Chl *d* and *f* (700-750 nm) in black, brown and pink beachrock samples (Fig. 1 – figure supplement 2). Spectral signatures of bacteriochlorophyll (BChl) *a* absorption (∼800 nm and ∼860 nm) indicative of the presence of anoxygenic phototrophic bacteria were found more scattered over the beachrock biofilm zonation. These findings are in line with previous, more detailed hyperspectral imaging studies of beachrock at Heron Island (Trampe and Kühl 2016, Kühl et al. 2020).

Pigment extraction and subsequent quantification of beachrock samples confirmed the presence of the above mentioned photopigments along with carotenoids and the cyanobacterial UV sunscreen pigment scytonemin (Garcia-Pichel et al. 1992) in the beachrock (Fig. 1). The scytonemin content per beachrock surface area was in the range of 20–130 mg scytonemin m^-2^, with highest amounts in black beachrock followed by brown and pink beachrock (Fig. 1A, B). This correlates with the air and UV exposure of the different beachrock zonations, where the black beachrock is found in the uppermost intertidal zone facing longer and more frequent periods of desiccation with strong solar radiation and thus UV exposure, while the brown and pink beachrock zones are found on lower parts of the beachrock platform subject to less desiccation and UV exposure (Cribb 1966; Trampe and Kühl, 2016). The scytonemin content of beachrock is similar to values found in densely pigmented hot spring microbial mat exposed to high UV exposure (Brenowitz and Castenholz, 1997; Dillon and Castenholz, 2003).

**Figure 1.**
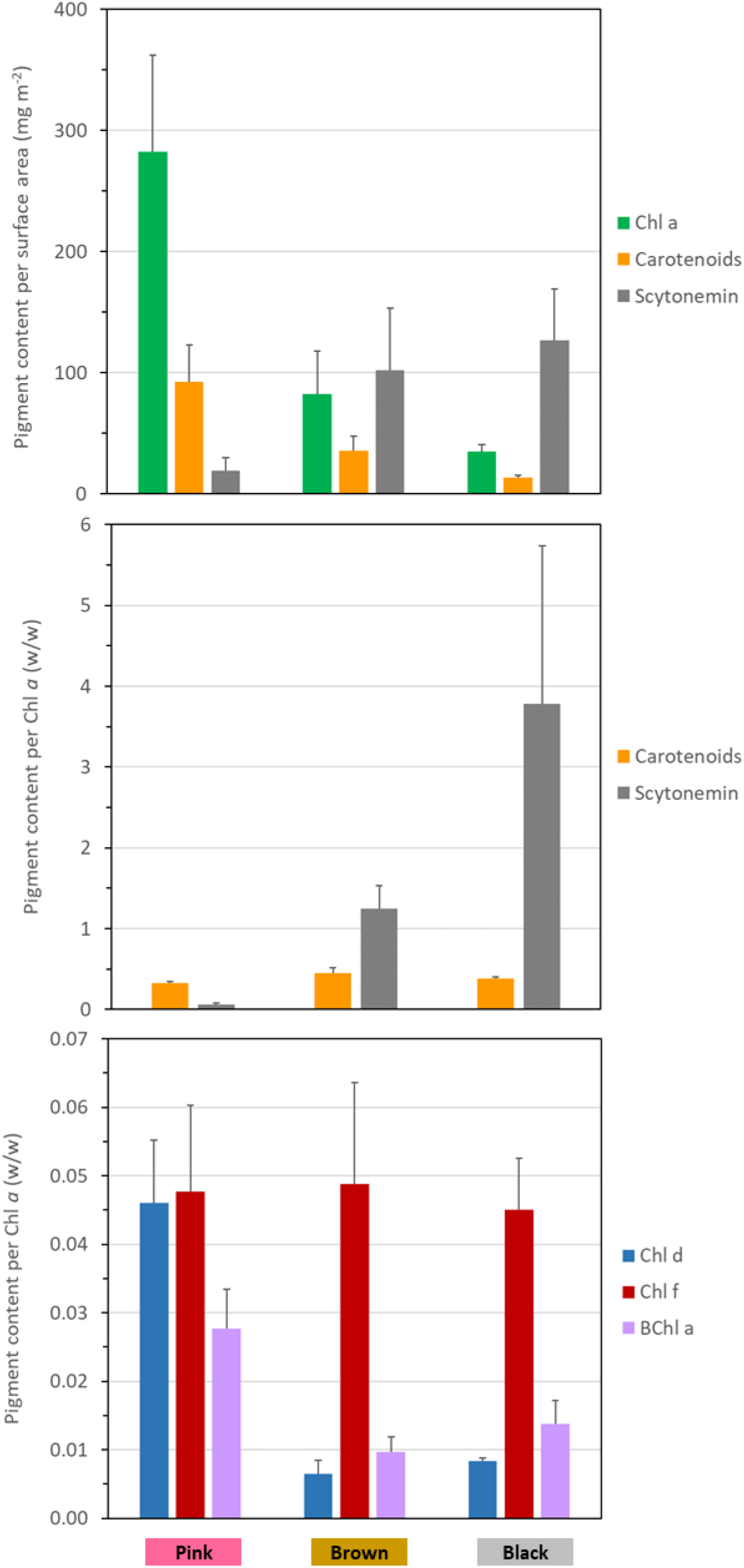
Quantification of chlorophyll, bacteriochlorophyll, carotenoids and the UV-protective pigment scytonemin in samples from three different beachrock zones (Bars ± error bars indicate means ± standard deviation; *n* = 3). Additional data are available in the Suppl. Materials showing color micrographs (Figure 1 - figure supplement 1) and hyperspectral images (Figure 1 – figure supplement 2) of beachrock cross-sections.

The Chl *a* content per beachrock surface area was in the range of 35–280 mg Chl *a* m^-2^ and the carotenoid content was in the range of ∼15-90 mg carotenoid m^-2^, with the highest content of both pigment types in pink beachrock followed by brown and black beachrock samples. The carotenoid to Chl *a* ratio was, however, similar (ca. 0.4 w/w) across all beachrock biofilm types (Fig. 1A), indicative of a pronounced photoprotective role of carotenoids for the beachrock microbial community.

Chl *f* was the second-most abundant Chl species in the beachrock after Chl *a*, and the Chl *f*-to-Chl *a* ratio was strikingly similar in all beachrock types (ca. 0.04–0.05 w/w relative to Chl *a*). Chl *d* and BChl *a* were found in variable amounts (0.01–0.05 w/w relative to Chl *a*), whereas the contents of Chl *b* and Chl *c* were very low or undetectable (<0.01 w/w relative to Chl *a*; data not shown). The coastal surface water contained Chl *a*, Chl *b*, and Chl *c* but no detectable amounts of other chlorophylls (data not shown).

### Mapping of O_2_ production in beachrock samples under visible and near infrared light

Imaging of beachrock cross-sections pressed against a planar optode in an experimental chamber (Fig. 2A) showed different O_2_ distributions in beachrock samples when illuminated vertically from above with predominantly visible (400-700 nm) or near-infrared (700-780 nm) light (Fig. 2B; Fig. 2 – figure supplement 1 and 2). Visible wavelengths and possibly a tail of NIR in the halogen lamp spectrum (Fig. 2 – figure supplement 3) stimulated O_2_ production in the upper biofilm at the beachrock surface, while no O_2_ production was observed in the endolithic zone. In contrast, illumination with NIR primarily stimulated O_2_ production in the endolithic, Chl *f*-containing layers. However, a small tail of VIS light in the spectrum of the 740 nm LED’s used for NIR illumination (amounting to a few % of the NIR photon irradiance; Table S1) lead to a weak stimulation of oxygenic photosynthesis in the surface biofilm. By introducing NIR long-pass filters in the light path (Fig. 2 – figure supplement 3) this small contribution of visible light was removed, albeit also at the cost of a lower NIR photon irradiance, and only O_2_ production in the endolithic layer remained (Fig. 2B; Fig. 2 – figure supplement 1 and 2). Such localized NIR-driven O_2_ production in the endolithic zone with Chl *f*-containing cyanobacteria was also observed when illuminating beachrock cross-sections homogeneously (Kühl et al., 2020), but in the present study we mapped the distribution of O_2_ production under more natural light conditions by illuminating the beachrock samples from above and thus generating a light gradient with depth in the sample, both with respect to photon irradiance and spectral composition. We conclude that NIR-driven photosynthesis in the beachrock is primarily taking place in the endolithic layer, where incident sunlight is strongly depleted in visible wavelength due to the dense overlaying biofilm at the beachrock surface. Such light microenvironment is conducive to induce FaRLiP in cyanobacteria (Gan et al., 2014; Trampe and Kühl, 2016), as the photoacclimation is regulated by phytochromes gauging the relative amount of visible and NIR (Zhao et al., 2015).

**Figure 2.**
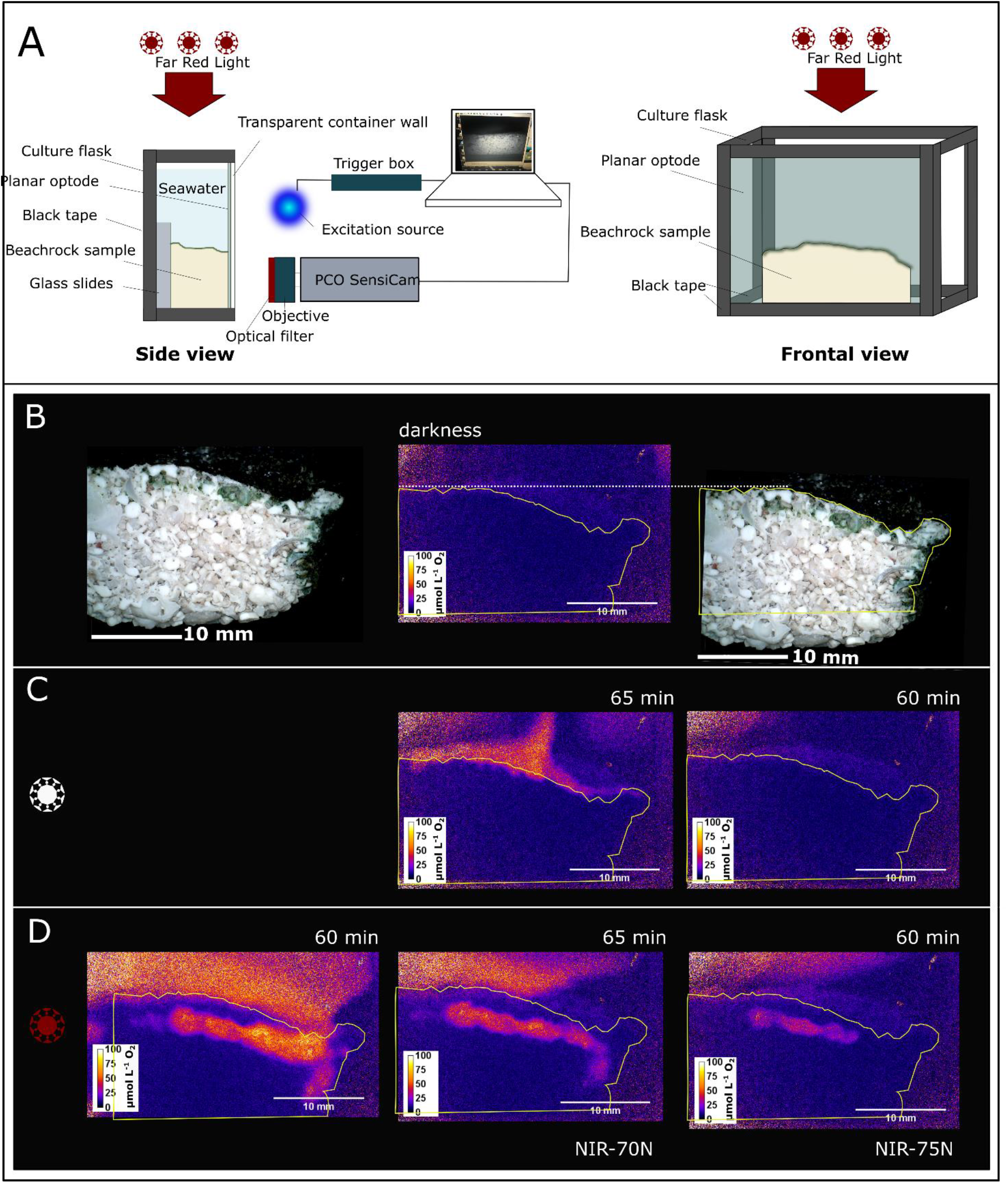
Luminescence lifetime imaging of dissolved oxygen on a black beachrock sample. (A) Experimental setup used for luminescence lifetime imaging of O_2_ in beachrock. Schematic drawing of the setup showing a side view of the chamber and the imaging system (left) and a frontal view of the experimental chamber (right). The beachrock sample is tightly pressed against the planar optode, using glass slides behind the sample, ensuring close contact. A computer controls the luminescence lifetime camera (PCO SensiCam SensiMod) used for read-out, as well as the trigger unit, triggering the blue LED (460 nm) exciting the luminescent dyes in the optode. All edges of the chamber are covered with black tape, to avoid scattering within the chamber walls. A calibrated NIR LED array or a halogen lamp was used to illuminate the beachrock surface from above with defined spectral photon irradiance levels. (B) Left: digital microscope image of the used sample. Middle: Outlined sample on a 2D image of O_2_ distribution (false color) acquired after equilibration in darkness. Right: Overlay of the sample outline with the digital microscope image. (C) Middle and Right: Color coded O_2_ distribution imaged after irradiation with white light [417 µmol photons m^-2^ s^-1^ (338_400-700nm_ + 79_700-780nm_ µmol photons m^-2^ s^-1^) and 86 µmol photons m^-2^ s^-1^ (70_400-700nm_ + 16_700-780nm_ µmol photons m^-2^ s^-1^)] for 65 minutes and 60 minutes, respectively. (D) Color coded O_2_ distribution imaged after illumination for 60, 65 and 60 minutes, respectively, with (left) a 740 nm LED unfiltered (1033 µmol photons m^-2^ s^-1^; 66_400-700nm_ + 967_700-780nm_ µmol photons m^-2^ s^-1^), (middle) a 740 nm LED + NIR-70N long pass filter (484 µmol photons m^-2^ s^-1^; 6_400-700nm_ + 478_700-780nm_ µmol photons m^-2^ s^-1^), and (right) a 740 nm LED + NIR-75N long pass filter (15 µmol photons m^-2^ s^-1^; 0.1_400-700nm_ + 15.0_700-780nm_ µmol photons m^-2^ s^-1^). Additional data for brown and pink beachrock sections are available in the Suppl. Materials (Figure 2 - figure supplements 1 and 2) along with spectra of the white and NIR light sources (Figure 2 – figure supplement 3) and a calibration curve of the planar optode (Figure 1 – figure supplement 4)

### Quantification of beachrock photosynthesis under visible and near infrared light

We used gas exchange measurements of net O_2_ fluxes under dark and varying light conditions to quantify the primary productivity of samples from the black, brown and pink zonation found on the beachrock platform on Heron Island (Fig. 3; Fig. 3 – figure supplement 1). Measurements of net O_2_ consumption and production overall showed that beachrock biofilms exhibit a high respiration activity both in dark and light conditions. Consequently, the compensation irradiance for visible light (400-700 nm), where beachrock switches from net consumption to net production of O_2_, was high reaching ∼100 µmol photons m^-2^ s^-1^ in brown beachrock, ∼200 µmol photons m^-2^ s^-1^ in pink beachrock, and ∼400 µmol photons m^-2^ s^-1^ in black beachrock (Fig. 3). The high respiration generally led to absence of net O_2_ production from beachrock incubated under NIR, with the exception of the brown biofilm, which showed a NIR compensation irradiance of ∼1000 µmol photons m^-2^ s^-1^. These measurements are in line with the O_2_ imaging results showing that NIR-induced O_2_ production in the endolithic part of the beachrock harboring cyanobacteria with FaRLiP was more or less completely consumed within the beachrock biofilm (Fig. 2). Nevertheless, when we calculated gross photosynthesis rates from the measured net gas fluxes, we found a substantial NIR-driven photosynthesis in the beachrock reaching >20% of the gross photosynthesis rates under comparable photon irradiances of visible light (Fig. 4). Comparing the initial quasi-linear slopes of the gross photosynthesis vs. photon irradiance curves, we found that the initial slopes of NIR-driven photosynthesis were 6, 8, and 10 % of the corresponding slopes for VIS-driven photosynthesis in pink, brown and black beachrock, respectively.

**Figure 3.**
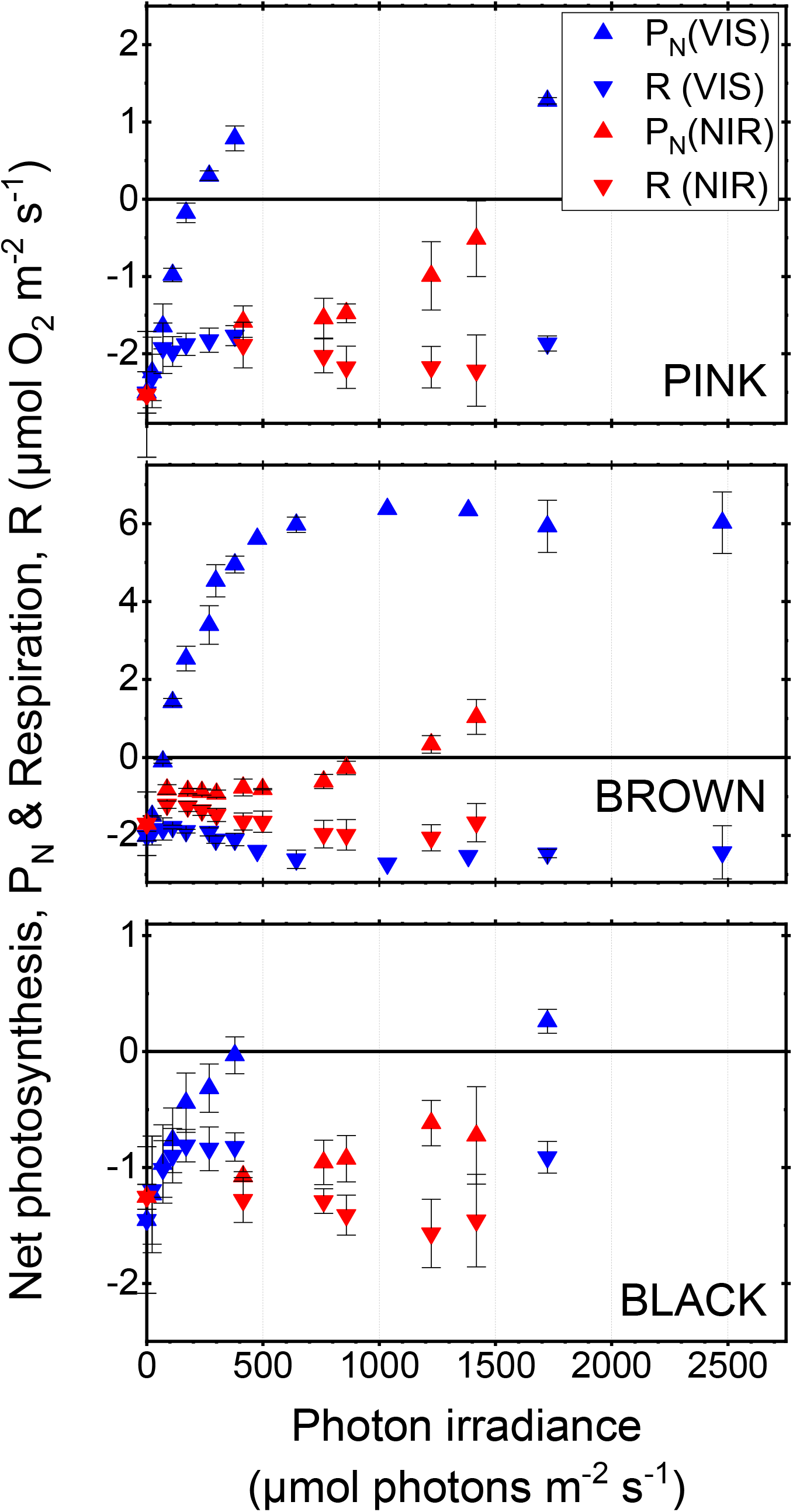
Gas exchange measurements of net photosynthesis (P_N_) and respiration (R) in samples from three different beachrock zones illuminated with visible light (VIS; 400-700 nm) and NIR (740 nm). Symbols ± error bars indicate means ± standard deviation; n=3). A schematic drawing of the experimental setup is available in the Suppl. Materials (Figure 3 - figure supplement 1).

**Figure 4.**
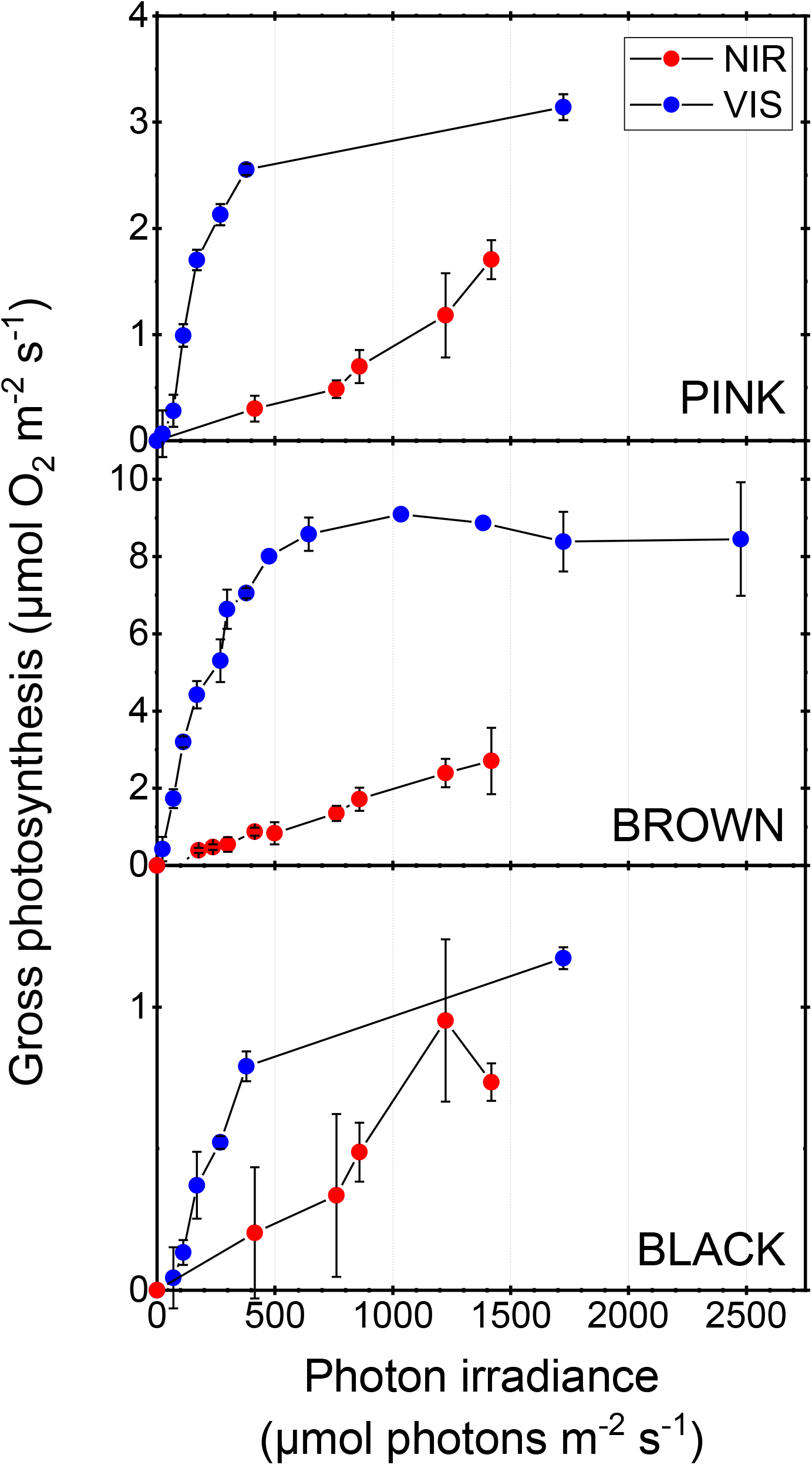
Areal gross photosynthesis rates in samples from three different beachrock zones illuminated with visible light (VIS; 400-700 nm) and NIR (740 nm). Symbols ± error bars indicate means ± standard deviation; n=3)

The VIS-driven gross photosynthesis in beachrock showed a typical saturation with increasing photon irradiance. Photosynthesis showed saturation at a high VIS photon irradiance of ∼500 µmol photons m^-2^ s^-1^ and exhibited no photoinhibition even at photon irradiances approaching the highest levels experienced in their natural habitat, i.e., ∼2000 µmol photons m^−2^ s^−1^. Such high light tolerance is probably linked to the large amounts of carotenoids and scytonemin in the beachrock biofilms (Fig. 1) acting as photoprotective pigments, as well as dynamic physiological adaptations such as non-photochemical quenching and redox balancing mechanisms (Schreiber et al., 2002; Kirilovsky and Kerfeld, 2016; Niyog and Truong, 2013; Muramatsu and Hihara, 2012).

The NIR-driven photosynthesis in the beachrock did not approach saturation and exhibited a continuously increasing rate with increasing photon irradiance (Fig. 4). This can be explained by the light attenuation due to scattering and absorption in the compacted beachrock biofilm, which prevented saturation of NIR-driven photosynthesis in the endolithic layer even at levels of incident light similar to solar irradiation on mid-day exposed beachrock. We were not able to precisely quantify the light reaching the endolithic layer in the different beachrock types when illuminated vertically from above, but a previous study (Kühl et al. 2020) showed that significant NIR-driven O_2_ production could be induced in the endolithic zone when illuminating a beachrock cross-section homogenously with low levels of NIR, i.e., 28 µmol photons m^−2^ s^−1^ (740 nm). NIR-driven photosynthesis in beachrock by cyanobacteria with FaRLiP thus appears relatively efficient. Recent studies of the photosynthetic apparatus of a cyanobacterium with FaRLip (*Chroococcidiopsis thermalis*) have shown that the inclusion of Chl *f* slows down energy trapping in the photosystems and can lead to a decreased efficiency of PSII (Mascoli et al., 2020). However, the combination of red-shifted phycobiliprotein antenna pigments with the modified photosystem configuration (by inclusion of Chl *d* and *f*) enlarges both the absorption and transfer of light to the photosystems and can speed up charge separation and enhance the quantum efficiency of PSII (Mascoli et al., 2022). Such capability of cyanobacteria with FaRLiP to dynamically fine tune their photosynthetic apparatus by switching between i) “normal” VIS-driven photosynthesis using phycobiliproteins and Chl *a*, and ii) NIR-driven photosynthesis using red-shifted phycobiliproteins and reaction centers modified with Chl *d* and *f*, thus seems a highly successful strategy in habitats like beachrock that experience strong fluctuations in irradiance and spectral composition of light due to tidal changes in water coverage.

Our measurements provide a first quantitative estimate of gross primary production in different types of beachrock, and also represent the very first estimate of the quantitative role of NIR-driven photosynthesis in a natural habitat harboring cyanobacteria with FaRLiP. While beachrock is a widespread sedimentary feature of intertidal (sub)tropical coasts (Vousdaukas et al., 2007), we are only aware of one published quantification of beachrock gross primary production amounting to ∼10 mmol O_2_ m^−2^ beachrock h^−1^ (Krumbein et al., 1979), as determined for water covered beachrock in an Egyptian sabkha habitat. In our study (see Figure 4), the maximal VIS-driven gross primary production ranged from ∼1 to 9 µmol O_2_ m^-2^ s^-1^, equivalent to 3.6 – 32.4 mmol O_2_ m^−2^ beachrock h^−1^, while maximal NIR-driven gross primary production ranged from ∼0.75 to 2.5 µmol O_2_ m^-2^ s^-1^, equivalent to 2.7 – 9 mmol O_2_ m^−2^ beachrock h^−1^. This span of gross photosynthesis rates falls well within the range of gross photosynthesis rates reported from other microbenthic biofilm habitats harboring microalgae and cyanobacteria (Krause-Jensen and Sand Jensen, 1994), further corroborating a potentially important role of cyanobacteria with FaRLip in natural habitats.

## SUMMARY

Beachrock surfaces are covered with compact photosynthetic biofilms, where extreme attenuation of visible light in the uppermost layer is partially compensated via NIR-driven photosynthesis in deeper layers harboring endolithic cyanobacteria expressing complementary photopigmentation via FaRLiP. Our data provide a more precise estimate of beachrock primary production under visible and near-infrared light, and we show that NIR-driven photosynthesis by cyanobacteria with FaRLiP can contribute significantly to benthic primary production, reaching up to ∼20% of visible light driven gross photosynthesis under similar photon irradiance. Such additional production points to an important but hitherto overlooked ecological role of cyanobacteria with FaRLiP in habitats (e.g. biofilms in shallow waters, terrestrial endolithic habitats and underneath dense terrestrial plant canopies), where visible light is more strongly attenuated than NIR. Given that cyanobacteria with FaRLiP capability have been found globally in many different environments, we argue that investigations of NIR-driven oxygenic photosynthesis are highly relevant in future studies of benthic primary production. Our study might also inspire the design of more efficient photobioreactors taking advantage of complementary spectral absorption based on cyanobacteria with FaRLiP, either alone or in combination with other oxygenic phototrophs that are primarily driven by visible light.

## MATERIALS AND METHODS

### Field site and beachrock sampling

We investigated samples from the upper black-brown zone and the pink zone of the intertidal beachrock platform on Heron Island (Great Barrier Reef, Queensland, Australia; 23°26.5540S, 151°54.7420E); the field site is described in detail elsewhere (Cribb, 1966; Díez et al., 2007; Trampe and Kühl, 2016). Beachrock fragments were sampled at low tide and cut into smaller subsamples with defined dimensions using a seawater-flushed band saw with diamond blade (DB-100; Inland Craft, USA) under dim light. These samples were subsequently stored dry and dark prior to experimental analyses at Heron Island Research Station, University of Queensland. Additional samples from the same beachrock zonations were cut in small pieces (∼1 x 1 x 1 cm^3^) and were frozen in liquid N_2_ on site and shipped on dry ice to Denmark for subsequent pigment extraction and HPLC analysis.

### Pigment analysis

Photopigments (chlorophylls and carotenoids) and the cyanobacterial UV screen scytonemin in the beachrock samples were quantified using a modified version of a previously described HPLC protocol (Trampe and Kühl, 2016; Kühl et al., 2020). Beachrock fragments with a defined surface area (ca. 3 cm^2^) and weight (ca. 4–7 g) were pulverized. A defined amount of pulverized beachrock (ca. 0.2 g) was extracted twice with acetone:methanol (7:2 by volume; each time 0.4 mL; pooled to give a total of 0.8 mL) using a sonicator as previously described (Kühl et al. 2020). The extract (0.8 mL) was cleared by centrifugation and the supernatant was filtered through a 0.2 μm pore-size syringe filter (Sartorius Minisart SRP 4 filter; Sartorius Ltd., Stonehouse, UK). The filtered extracts were loaded onto a cooled HPLC autosampler (Agilent Technologies, Santa Clara, CA) and analyzed within 8 hours. The injected volume was 100 μL. The HPLC setup was similar to previously described setup (Kühl et al. 2020), except that the solvent system was modified as follows: The gradient of solvent A (methanol:acetonitrile:water, 42:33:25 by volume, supplemented with 10 mM ammonium acetate), and solvent B (methanol:acetonitrile:ethylacetate, 50:20:30 by volume) was changed linearly from 0% solvent B at the time of injection to 50% at 15 min, to 100% at 52 min, staying at 100% for 15 min before returning to 0% in 1 min. The chromatograph peaks were identified by comparison of their retention time and absorption spectrum with published data (Garcia-Pichel et al. 1992; Frigaard et al. 1996; Kühl et al. 2020). Pigments were quantified from the integrated peak area and using published absorption coefficients for chlorophylls and carotenoid in methanol (Garcia-Pichel et al. 1992; Frigaard et al. 1996; Li et al. 2012). This method of pigment quantification by HPLC was validated by injecting 100 μL of methanol with a known concentration of pure Chl *a* (*A*_665_ = 0.54; cat. no. 96145; Supelco, US) into the HPLC.

The calculated amount of Chl *a* obtained by HPLC analysis using peak integration was 93% of the Chl *a* amount calculated from the spectrum in methanol. Some of the deviation from 100% recovery could be due to a difference in absorption coefficient of Chl *<u>a</u>* in methanol and in the HPLC solvent at the time of elution. For sake of simplicity and a lack of pigment standards other than Chl *a*, the absorption coefficients used for quantification of all pigments by HPLC were assumed to be identical to that in methanol. The amount of pigments per surface area of beachrock was calculated from the amount of pigments determined by HPLC and the beachrock surface area represented by the amount of beachrock extracted (see above). All pigment contents are reported as the average and standard deviation of three separate extractions and HPLC analyses on pulverized sub-samples from the same beachrock zonation samples used for gas exchange measurements.

### Hyperspectral imaging and microscopy

The distribution of photopigments over beachrock cross-sections was mapped with a hyperspectral imaging system (Snapscan VNIR; imec-int.com) equipped with a color-corrected objective (Apo-Xenoplan, 2.0/20-0003; Schneider-Kreuznach GmbH, Germany). The camera system records hyperspectral image cubes by moving a filter plate in front of the camera chip (Geelen et al., 2013), enabling hyperspectral imaging of reflectance or luminescence with up to 150 bands/channels covering a wavelength range from ∼470 to 900 nm at high spatial resolution (7Mpx per spectral band). A more detailed description of the imaging system can be found in Zieger et al. (2020). Hyperspectral imaging was done on beachrock cross sections that were placed in a glass petri dish and covered with a thin (∼3-5 mm) layer of seawater. The Petri dish was placed directly below the camera system, which was mounted vertically above. For hyperspectral reflectance imaging, the beachrock samples were illuminated homogeneously with 4 halogen lamps (OSRAM DECOSTAR®; 20W, 205 lm; Osram, Germany) mounted at a 45° angle relative to the petri dish. Prior to measurements of reflected light from the beachrock sample, we acquired a hyperspectral image stack of reflected light from a white 95% reflectance target (T95; imec-int.com) placed at the same distance and position in the light field as the beachrock samples. Acquired hyperspectral image stacks of reflected light from beachrock samples were normalized with the hyperspectral image stack of reflected light from the reflectance standard yielding hyperspectral image stack of %reflectance. Due to a built-in shutter, all acquired images were automatically corrected for dark noise in the camera system. Image acquisition and subsequent image correction and analysis was done using the camera system software (HIS Studio; imec-int.com).

Color RGB images of the scanned beachrock cross-sections were generated by the system software, selecting spectral bands in the red (602.8 nm), green (536.4 nm) and blue (472.9 nm) part of the spectrum, to provide close to true color rendering. Subsequently, the software was used to find and highlight areas with similar spectral patterns. Using the Imec Spectral Angle Classifier in the system software, we could highlight areas with identical spectral properties over the coral cross sections. For this, we trained the classifier by defining small areas of the cross section with particular spectral features (reflectance spectra with and without signatures of Chl *f*), where each selection was treated as a separate class by the classifier. The classifier analysis yielded false color-coded maps of the coral cross-sections highlighting regions with similar spectral properties.

Color images of beachrock sections were also obtained with a digital microscope mounted on a stand with a focus drive (DinoLite Edge 3.0; Dinolite.com) using the built-in LED light of the microscope head. Camera control and image acquisition was done with the manufacturer’s software (DinoCapture 2.0 Version 1.5.37).

### Oxygen imaging

The depth distribution and dynamics of O_2_ in beachrock samples under VIS and NIR irradiance, was quantified by luminescence lifetime imaging using an O_2_-sensitive planar optode that was pressed tightly against a smooth beachrock cross-section (Fig. 2 – figure supplement 1). The fabrication of planar optodes and experimental set-ups for such measurements have been described in detail elsewhere (e.g. Kühl et al. 2008; Koren and Kühl 2018; Santner et al. 2015; Mosshammer et al. 2019). Planar optodes for imaging dissolved O_2_ distributions were made out of the O_2_-sensitive indicator dye platinum(II)meso-tetra(4-fluorophenyl)tetrabenzoporphyrin (Pt-TPTBPF4, 0.3% w/v) generously provided by Dr. Sergey Borisov (Institute for Analytical Chemistry and Food Chemistry, Graz University of Technology) and the antenna dye Fluoreszenzrot (FR, 0.3% w/v) purchased from Kremer Pigments (www.kremerpigments.com), both dissolved in a cocktail of polystyrene (PS) in toluene (10% w/v). The sensor cocktail was coated in a thin layer onto a transparent carrier foil (PET, GoodFellow) using a knife coater (www.byk-instruments.com). After solvent evaporation the sensor-layer was ∼ 12 µm thick. Further details on the fabrication are given elsewhere (Mosshammer et al. 2019)

A modular, time-domain-based luminescence lifetime imaging system was used for the O_2_ imaging as described in detail elsewhere (Holst et al., 1998; Kühl et al. 2008). A modulated CCD camera (SensiMod, PCO, Germany) with a camera lens (Xenoplan 1.4/17 CCTV-Lens, Schneider-Kreuznach, Germany) mounted with a 695 nm long pass filter (O92 Dunkelrot, Schneider-Kreuznach, Germany) was used to image the O_2_-dependent luminescence lifetime of the indicator (max emission at ∼780 nm), while filtering out background luminescence e.g. from fast-decaying chlorophyll fluorescence. We used a 460 nm LED (Photonic Research Systems Ltd, UK) for excitation of the luminescent dyes for a defined time interval (40 µs), while two intensity images (A_1_ and A_2_) of the resulting luminescence were recorded over 2 time windows (10 µs each) after the LED pulse (W_1_ at t_1_ = 41 µs, and W_2_ at t_2_ = 52 µs). From such pairs of dark-corrected luminescence intensity, the luminescence lifetime (τ) was calculated as (Equation 1; Holst et al. 1998):

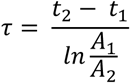

Image acquisition was controlled by the system software (look@molli; Holst and Grunwald, 2001) and lifetime calculation was done in the freely available image analysis software ImageJ in Fiji (Schindelin et al. 2012). All luminescence lifetime imaging was done in filtered seawater (salinity 35) in a temperature controlled dark room kept at 26°C.

For imaging, a planar optode (7 X 10 cm) was mounted on the inside of a plastic chamber, i.e., a cut 200 mL culture flask, with transparent walls (Fig.2 – figure supplement 1). The bottom of the planar optode was glued first, then a droplet of water was added between the container wall and the optode. The planar optode was placed on the water droplet, carefully smoothing it out and removing any trapped air bubbles. Then the other three sides were sealed using black tape (see more details in Mosshammer et al 2019). Black tape was also used to cover all edges, and sides (excluding the side with the planar optode) of the chamber to minimize light scattering through the container walls.

The chamber was illuminated from above using either a fiber-optic halogen lamp equipped with a collimating lens (KL2500LCD, Schott, Germany) or an array of NIR LEDs (740 nm; LZ4-40R300, LED Engin, Inc, San Jose, CA; HBW 25 nm) in combination with different longpass filters (Clarex NIR-70N and NIR-75N purchased from Weatherall). Black tape was used to define the window of illumination, and avoid light reaching the cut sides of the beachrock sample. Incident photon irradiance levels for different lamp or LED settings were quantified with a calibrated spectroradiometer (BTS256, Gigahertz Optics GmbH; 1 nm spectral resolution, spectral range 360-830 nm; Fig. 2 – figure supplement 2, Table S1).The mounted O_2_ planar optode was calibrated in the chamber using N_2_ gas and a calibrated reference sensor (OXR430-OI, Pyroscience GmbH) connected to a fiber-optic O_2_ meter (FireSting GO_2_, PyroScience GmbH). The air-saturated seawater in the chamber was flushed for some time with N_2_ and the chamber was then closed with a lid. After the N_2_ flow was stopped and once the reference sensor reached a stable value, indicating a steady state regarding dissolved O_2_ in the chamber, a calibration image was acquired. This was done repeatedly at declining O_2_ concentrations to obtain a calibration curve (Fig. 2 –figure supplement 3). The zero O_2_ value was reached chemically, by adding Na_2_SO_3_ to the chamber. After calibration, the chamber was cleaned to remove any Na_2_SO_3_ residue prior to the experiments.

Calibration values of the luminescence lifetime (averaged over the planar optode area) at defined O_2_ concentrations were fitted with an exponential decay function (R^2^=0.997), and could also be described well by a Stern-Volmer plot using the two-site model (R^2^=0.999; Klimant et al. 1997) (Equation 2):

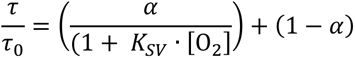

where τ_0_ is the sensor luminescence lifetime at zero O_2_, τ is the luminescence at a given O_2_ concentration, K_SV_ is the quenching constant of the sensor material, and α is the quenchable fraction of the immobilized sensor dye in the planar optode. This enabled conversion of acquired luminescence lifetime images during experiments to concentration images describing the 2D O_2_ distribution over the beachrock cross-sections.

For the sample measurements, smooth beachrock cross-sections were pressed tightly against the planar optode ensuring good contact and no water circulation between the beachrock cross section and the planar optode. Microscope slides were put behind the beachrock sample to push it again the planar optode. The seawater in the chamber was exchanged for every measurement series and the sample was given time to equilibrate in darkness for ca. 30 min prior to measurements under defined irradiance levels of VIS or NIR. All measurements were done in stagnant water to further minimize any potential effect of water circulation between sample and optode.

### Gas exchange measurements

Areal rates of net photosynthesis and respiration were measured at 26°C as net fluxes of O_2_ on ∼1-1.5 cm thick cylindrical beachrock samples with a defined surface area (3 technical replicate measurements on a samples from each beachrock zone). The beachrock samples were trimmed and carefully covered with black tape on the sides to fit tightly inside a small gas-exchange chamber made of transparent perspex (Fig. 3 – figure supplement 1). The chamber contained a small magnetic stir bar and an optical O_2_ sensor spot with a black optical isolation (OXSP5, Pyroscience GmbH), which was mounted on the inside and read out via a fiber-optic cable (SPFIB-BARE, Pyroscience GmbH) fixed to the chamber wall at one end and connected to a fiber-optic O_2_ meter (FireStingO_2_, Pyroscience GmbH) at the other end.

The gas exchange chamber was positioned next to a magnetic stirrer and was illuminated either with a white LED lamp (KL2500 LED, Schott GmbH) or an array of near infrared LEDs (740 nm; LZ4-40R300, LED Engin, Inc, San Jose, CA; HBW 25 nm) controlled via a custom-built driver. Defined levels of white light were adjusted by settings on the LED lamp, while different levels of NIR were adjusted by varying the number of NIR LEDs and their distance to the gas-exchange chamber. The downwelling irradiance was quantified for different light settings of NIR or white light illumination with a calibrated spectroradiometer (BTS256, Gigahertz Optics GmbH; 1 nm spectral resolution, spectral range 360-830 nm) positioned at the level of the beachrock sample surface in the experimental chamber.

Spectra of downwelling irradiance (acquired in units of µW m^-2^ nm^-1^), were converted to spectra of downwelling photon irradiance (in units of µmol photons m^-2^ s^-1^ nm^-1^) using Planck’s equation: E_λ_=h·c/λ, where E_λ_ is the energy of a photon with wavelength, λ, h is Planck’s constant (6.626 × 10^−34^ W s^2^), and c is the speed of light in vacuum (in m s^−1^). Subsequently, the integral photon irradiance over defined spectral regions (UV = 360-400 nm, VIS = 400-700 nm and NIR = 700-830 nm) was found by simple integration.

After mounting of a beachrock sample, the gas-exchange chamber was filled with aerated seawater (S=35, T=26°C) and closed tightly, while taking care no gas bubbles were trapped in the chamber. The chamber was positioned under the light source, and the change in O_2_ concentration was logged over time in darkness and for defined irradiance levels. Due to a relative small water volume (35-38 mL) relative to the beachrock sample surface area (∼6.2 cm^2^), a constant O_2_ depletion or production rate for a given light setting could be detected as a linear change in O_2_ concentration over 5-10 min, as determined by linear regression on parts of the O_2_ concentration vs. time measurements. After measurements, all samples were frozen in liquid N_2_ on site and shipped on dry ice to Denmark for subsequent pigment extraction and HPLC analysis (see above).

The net flux of O_2_ was calculated from the measured O_2_ concentration depletion or production rate (dC/dt) as J = (V/A)·dC/dt, where V is the volume of water in the chamber above the beachrock, and A is the surface area of the beachrock sample. Dark respiration, R_D_, was measured as the net O_2_ flux in dark incubated samples, while net photosynthesis P_N_ (= P_G_ – R_L_) was measured as the net O_2_ flux in light incubated samples, where P_G_ is the gross photosynthesis and R_L_ is the respiration in light. We estimated R_L_ from measurement of post-illumination respiration, i.e., by measurements of the net O_2_ consumption rate in the dark immediately after a period with illumination at each photon irradiance level (Cooper et al., 2011). The gross photosynthesis at each photon irradiance level was estimated as P_G_ = P_N_ + R_L_.

## ACKNOWLEDGEMENTS

This study was supported by grants from the Independent Research Fund Denmark (MK; DFF-8022-00301B & DFF-8021-00308B), an investigator award from the Gordon and Betty Moore Foundation (MK; grant no. GBMF9206; https://doi.org/10.37807/GBMF9206), and an Experiment grant from the Villum Foundation (MM; VIL 50371). We thank Sofie Jakobsen, Veronica Pedersen, Victoria Thuesen and Mary Rose Gonzalez for excellent technical support and sample analyses. We thank the staff of Heron Island Research Station, University of Queensland, for technical and logistic support. Work at Heron Island was conducted under permit no. G16/38423.1 and G22/46725.1 from the Great Barrier Reef Marine Parks authority.

## AUTHOR CONTRIBUTIONS

Maria Mosshammer: Conceptualization, Investigation, Formal analysis, Data curation, Validation, Visualization, Methodology, Writing, Review and editing.

Erik Trampe: Conceptualization, Investigation, Formal analysis, Data curation, Visualization, Methodology, Review and editing.

Niels Ulrik Frigaard: Conceptualization, Resources, Funding acquisition, Data curation, Investigation, Formal analysis, Methodology, Review and editing.

Michael Kühl: Conceptualization, Resources, Formal analysis, Supervision, Funding acquisition, Validation, Investigation, Visualization, Methodology, Project administration, Writing, Original draft, Review and editing.

---

**Figure 1 – figure supplement 1.**
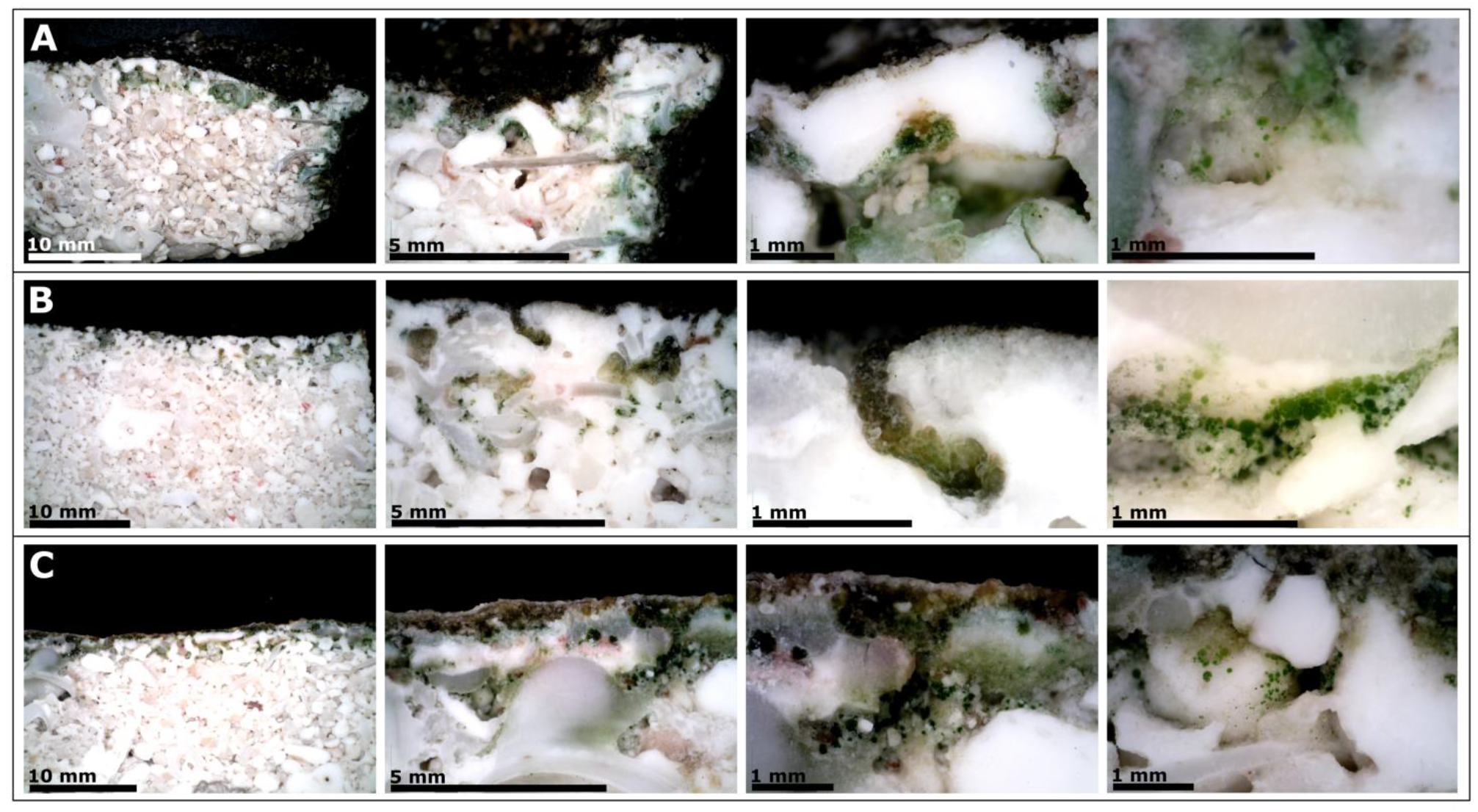
Beachrock samples used for O_2_ imaging. Digital microscope images of black (A), brown (B) and pink (C) beachrock samples at various magnifications (see scale bar in each panel). The microscope images were taken after completion of the O_2_ imaging.

**Figure 1 – figure supplement 2.**
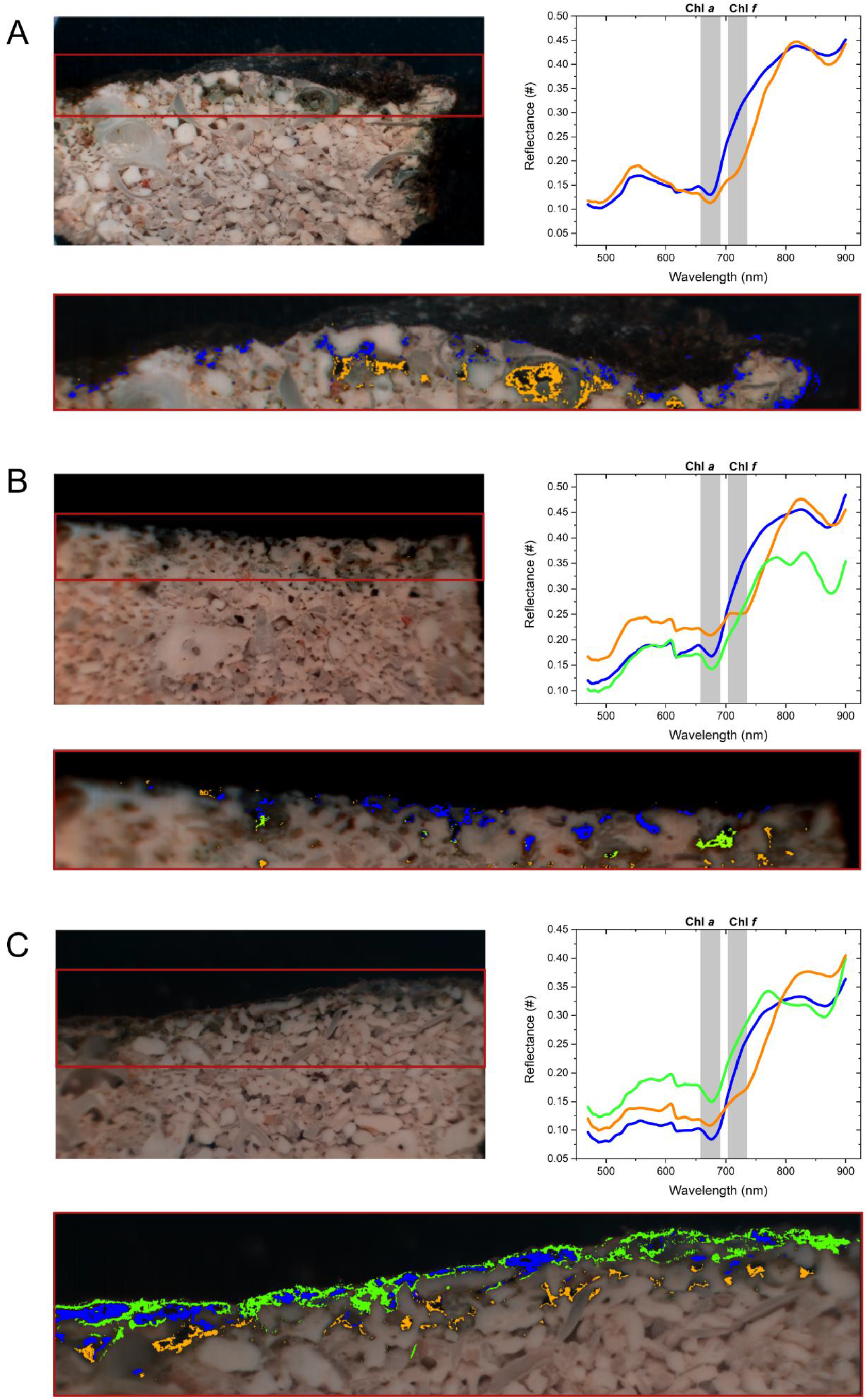
Hyperspectral reflectance imaging of beachrock samples. Left: Black (A), brown (B) and pink (C) beachrock samples with and without selected regions of interest (color coded) of similar spectral composition. Right: Spectral information retrieved from the selected regions. Hyperspectral reflectance measurements were conducted subsequently to the O_2_ mapping experiments.

**Figure 2 – figure supplement 1.**
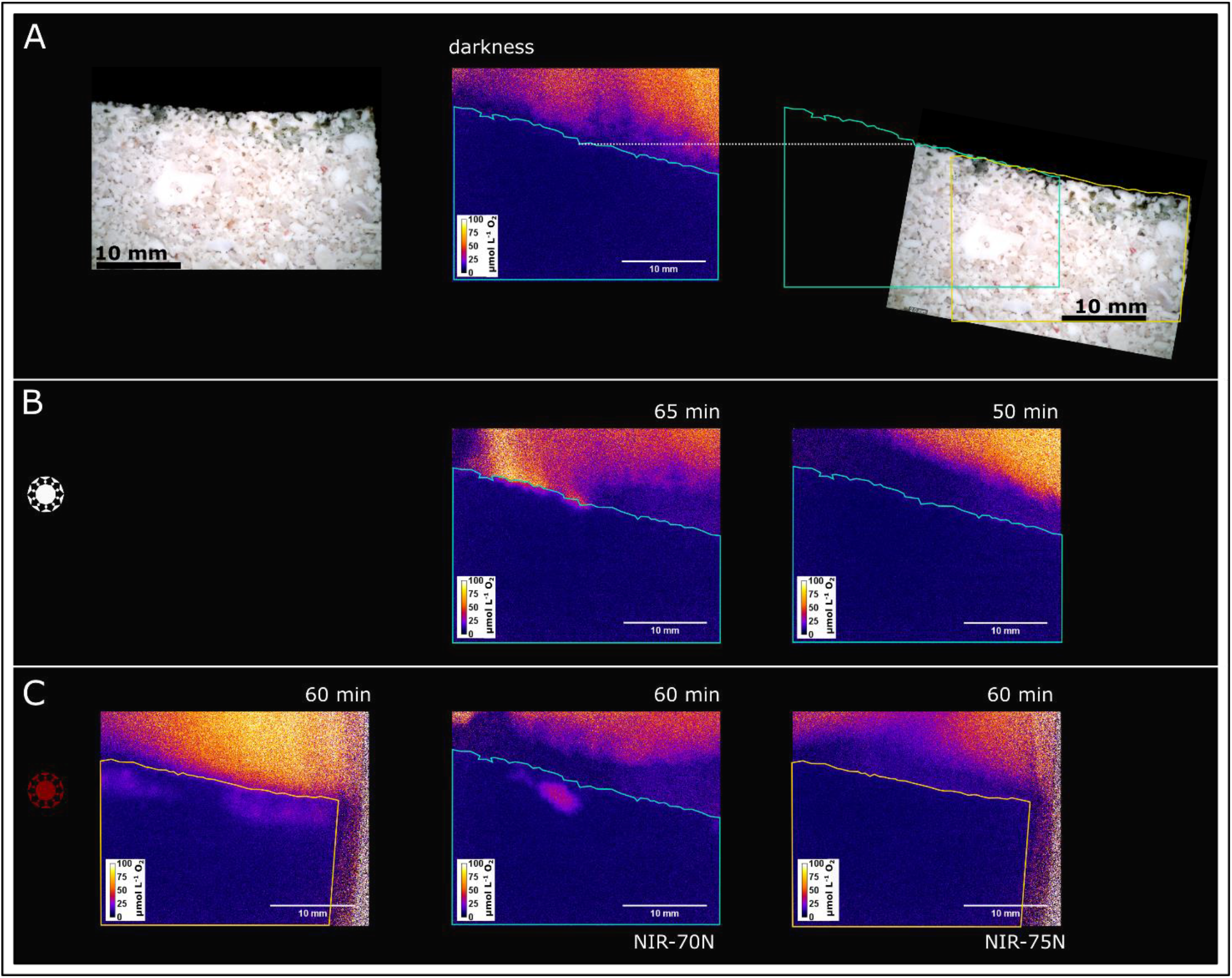
Luminescence lifetime imaging of dissolved oxygen on a brown beachrock sample. (A) Left: digital microscope image of the used sample. Middle: Outlined sample on a 2D image of O_2_ distribution (false color) acquired after equilibration in darkness. Right: Overlay of the sample outline with the digital microscope image. (B) Middle and Right: Color coded O_2_ distribution imaged after irradiation with white light (417 µmol photons m^-2^ s^-1^ (338_400-700nm_ + 79_700-780nm_ µmol photons m^-2^ s^-1^) and 86 µmol photons m^-2^ s^-1^ (70_400-700nm_ + 16_700-780nm_ µmol photons m^-2^ s^-1^)) for 65 minutes and 50 minutes, respectively. (C) 2D images of dissolved O_2_ acquired after 60 minutes irradiation with (left) a 740 nm LED unfiltered (131 µmol photons m^-2^ s^-1^ ; 9_400-700nm_ + 122_700-780nm_ µmol photons m^-2^ s^-1^), (middle) a 740 nm LED + NIR-70N long pass filter (484 µmol photons m^-2^ s^-1^; 6_400-700nm_ + 478_700-780nm_ µmol photons m^-2^ s^-1^), and (right) a 740 nm LED + NIR-75N long pass filter (22 µmol photons m^-2^ s^-1^; 0.1_400-700nm_ + 22_700-780nm_ µmol photons m^-2^ s^-1^).

**Figure 2 – figure supplement 2.**
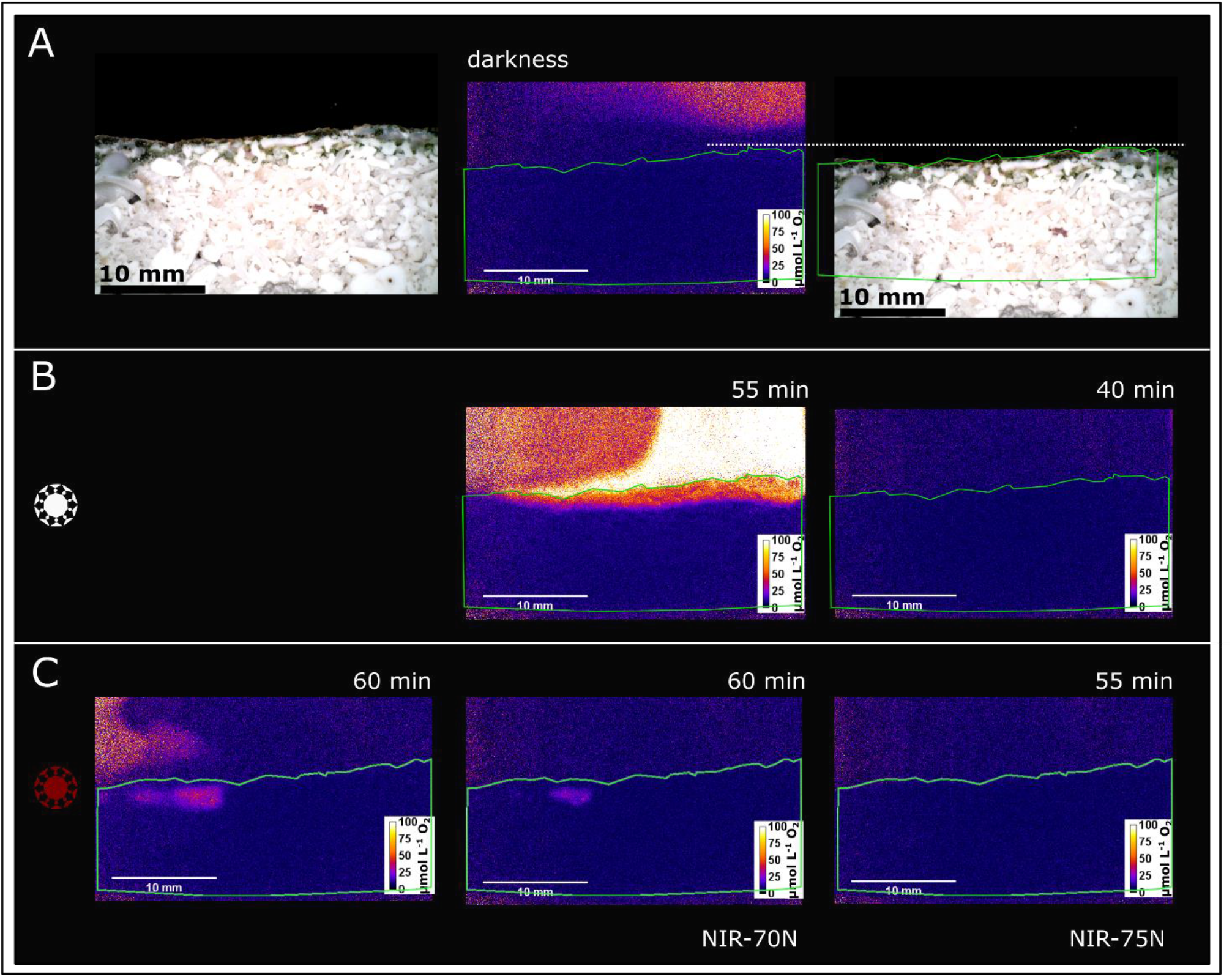
Luminescence lifetime imaging of dissolved oxygen on a pink beachrock **sample.** (A) Right: digital microscope image of the used sample. Middle: Outlined sample on a 2D image of O_2_ distribution (false color) acquired after incubation in darkness. Left: Overlay of the sample outline with the digital microscope image. (B) Middle and Right: Imaged O_2_ distribution, after irradiation with white light [417 µmol photons m^-2^ s^-1^ (338_400-700nm_ + 79_700-780nm_ µmol photons m^-2^ s^-1^) and 86 µmol photons m^-2^ s^-1^ (70_400-700nm_ + 16_700-780nm_ µmol photons m^-2^ s^-1^)] for 55 minutes and 40 minutes, respectively. (C) Imaged O_2_ distribution after 60, 60 and 55 minutes illumination, respectively, with (left) a 740 nm LED unfiltered (1033 µmol photons m^-2^ s^-1^;66_400-700nm_ + 967_700-780nm_ µmol photons m^-2^ s^-1^), (middle) a 740 nm LED + NIR-70N long pass filter (484 µmol photons m^-2^ s^-1^; 6_400-700nm_ + 478_700-780nm_ µmol photons m^-2^ s^-1^), and (right) a 740 nm LED + NIR-75N long pass filter, 15 µmol photons m^-2^ s^-1^ (0.1_400-700nm_ + 15_700-780nm_ µmol photons m^-2^ s^-1^).

**Figure 2 – figure supplement 3.**
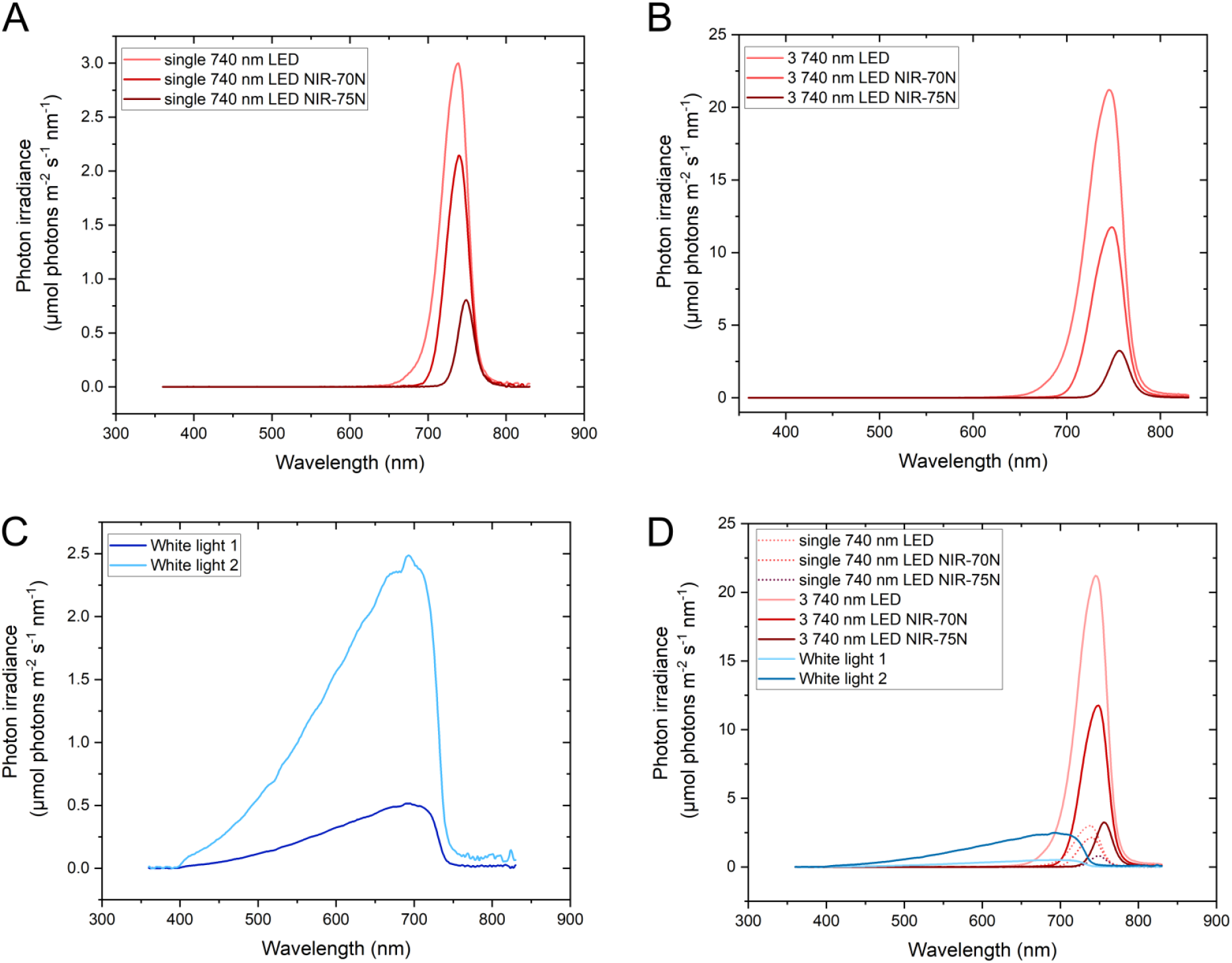
Spectra of the light sources used in the O_2_ imaging experiments. A) Single 740 nm LED with and without NIR-70N and NIR-75N longpass filter; B) three 740 nm LEDs LED with and without NIR-70N and NIR-75N longpass filter; C) used white halogen lamp (Schott KL2500 LCD), D) all combined.

**Figure 2 – figure supplement 4.**
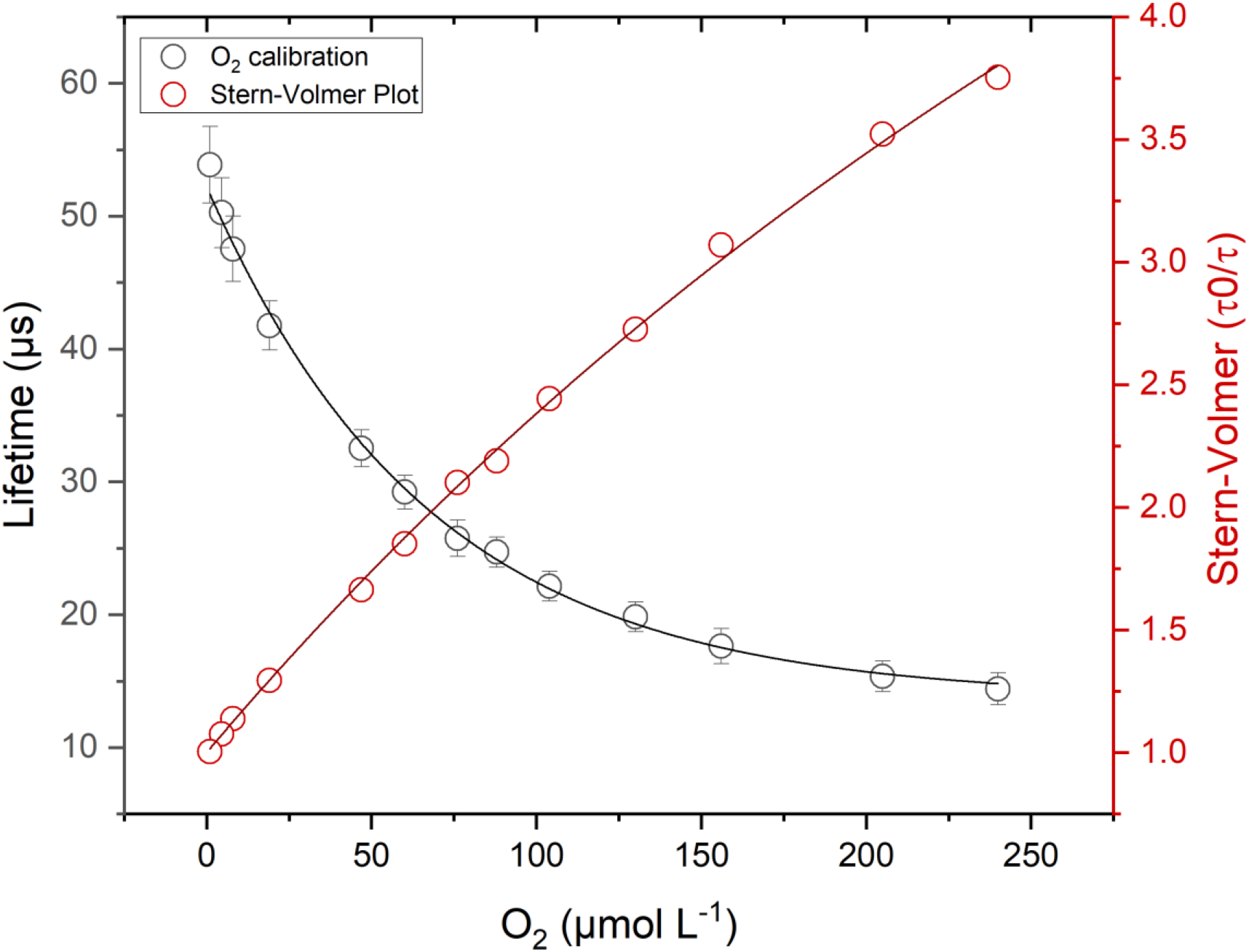
Calibration curve of the O_2_ planar optode and the correlating Stern-Volmer Plot. The calibration curve was fitted with an exponential decay function (R^2^=0.997) and the Stern-Volmer Plot with the simplified Two-Site Model (Eq. 2) (R^2^=0.999). Symbols and error bars represent means ± standard deviation of pixel values across the imaged part of planar optode.

**Figure 3 – figure supplement 1.**
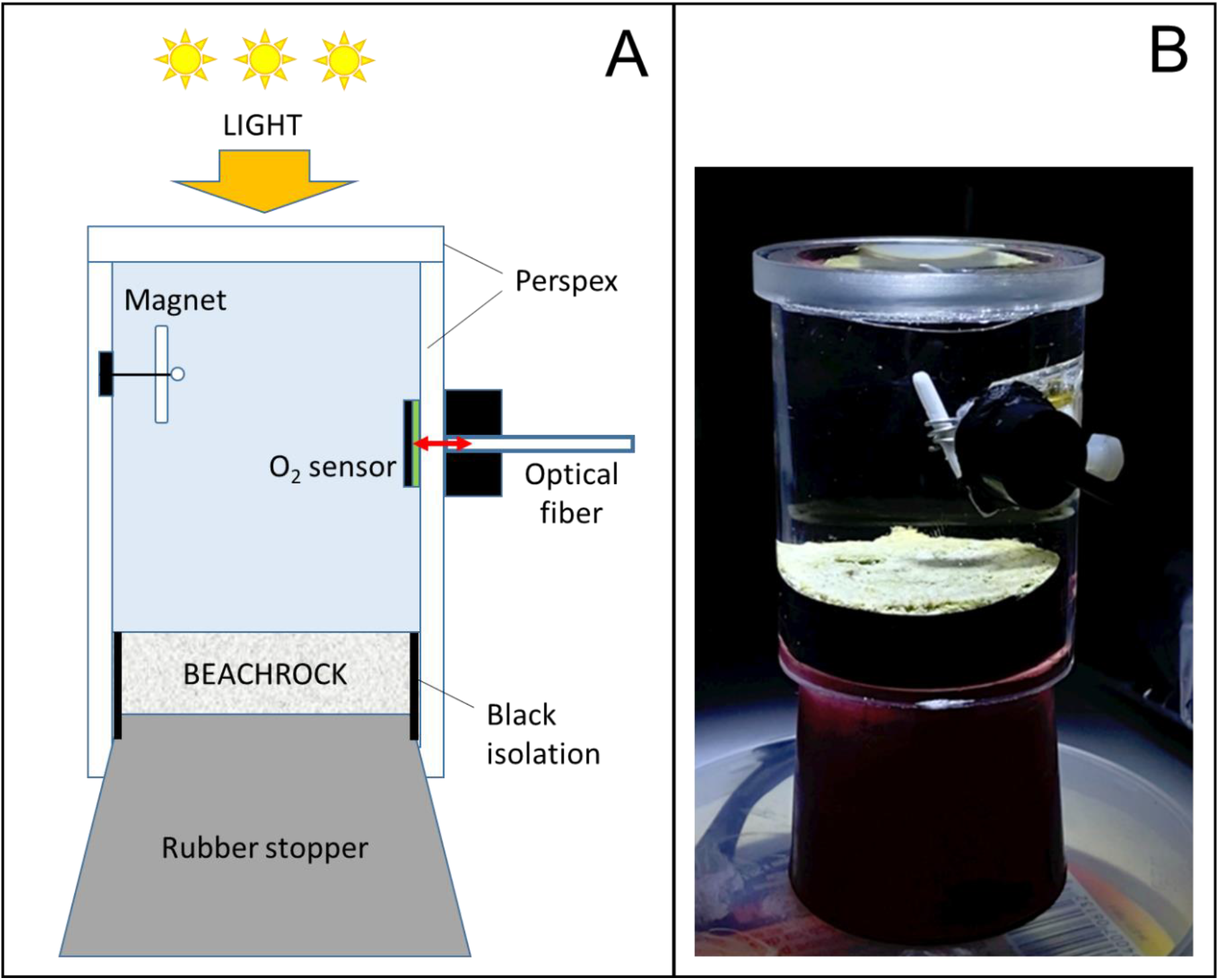
Experimental chamber used for gas-exchange measurements on beachrock. (A) Schematic drawing of experimental setup. (B) Photo of beachrock sample mounted in gas-exchange chamber. Note that the sides of the sample are shielded from light exposure by black tape. The O_2_ concentration in the chamber is monitored with an optical O_2_ sensor patch mounted on the inside and read-out by an optical fiber, which is mounted on the outside of the transparent Perspex chamber wall and connected to a fiber-optic O_2_ meter. The sensor patch was covered by a black O_2_ permeable optical isolation to avoid interference from background light on the sensor signal. Stirring in the chamber is induced by a small magnet mounted on the chamber wall and driven by a magnetic stirrer positioned next to the chamber. The chamber was illuminated from above with LED light of known intensity and spectral composition.

**Table S1.** Incident irradiance levels of white and NIR light used for illumination during 2D mapping of dissolved O_2_ concentrations in beachrock.

| Light source<br>+ filter | 360 – 830 nm<br>($\mu\text{W m}^{-2} \text{ nm}^{-1}$) | 400 – 700 nm<br>($\mu\text{mol photons m}^{-2} \text{ s}^{-1}$) | 700-780 nm<br>($\mu\text{mol photons m}^{-2} \text{ s}^{-1}$) | 400 – 780 nm<br>($\mu\text{mol photons m}^{-2} \text{ s}^{-1}$) |
| --- | --- | --- | --- | --- |
| 740 nm (1 LED) | 21.41 | 9.15 | 121.55 | 130.7 |
| 740 nm (1 LED)<br>+ NIR-70N filter | 12.65 | 1.01 | 76.70 | 77.71 |
| 740 nm (1 LED)<br>+ NIR-75N filter | 3.54 | 0.09 | 22.04 | 22.13 |
| 740 nm (3 LEDs) | 168.59 | 65.51 | 967.43 | 1032.94 |
| 740 nm (3 LEDs)<br>+ NIR-70N filter | 78.43 | 5.68 | 478.14 | 483.83 |
| 740 nm (3 LEDs)<br>+ NIR-75N filter | 15.23 | 0.10 | 14.96 | 15.06 |
| White halogen light setting 1 | 16.63 | 69.83 | 16.61 | 86.44 |
| White halogen light setting 2 | 80.30 | 337.79 | 79.32 | 417.11 |

## REFERENCES

Antonaru, L.A., Cardona, T., Larkum, A.W.D. et al. (2020) Global distribution of a chlorophyll *f* cyanobacterial marker. ISME J 14: 2275–2287.

Billi D, Napoli A, Mosca C, Fagliarone C, de Carolis R, Balbi A, Scanu M, Selinger VM, Antonaru LA, and Nürnberg DJ (2022) Identification of far-red light acclimation in an endolithic *Chroococcidiopsis* strain and associated genomic features: Implications for oxygenic photosynthesis on exoplanets. Front Microbiol 13: 933404.

Brenowitz, S., and Castenholz, R. W. (1997) Long-term effects of UV and visible irradiance on natural populations of a scytonemin-containing cyanobacterium (*Calothrix* sp.). FEMS Microbiol Ecol 24: 343–352, 10.1111/j

Chen M, Schliep M, Willows R, Cai Z-L, Neilan B, and Scheer H (2010) A red-shifted chlorophyll. Science 329: 1318–1319.

Chen M, and Blankenship RE (2011). Expanding the solar spectrum used by photosynthesis. Trends Plant Sci 16:427–31.

Chen M, Li Y, Birch D, and Willows RD (2012) A cyanobacterium that contains chlorophyll *f* — a red-absorbing photopigment. FEBS Lett, 586: 3249–3254.

Chen M (2014) Chlorophyll modifications and their spectral extension in oxygenic photosynthesis. Ann Rev Biochem 83: 317–340.

Cooper, T. F., Ulstrup, K. E., Dandan, S. S., Heyward, A., Kühl, M., Muirhead, A., O’Leary, R., Ziersen, B and van Oppen, M. J. H. (2011). Niche specialisation of reef-building corals in the mesophotic zone: metabolic trade-offs between divergent *Symbiodinium* types. Proc Roy Soc London: B 278: 1840–1850.

Diez B, Bauer K, and Bergman B (2007) Epilithic cyanobacterial communities of a marine tropical beach rock (Heron Island, Great Barrier Reef): diversity and diazotrophy. Appl Environ Microbiol 73: 3656–68.

Dillon, J. G., and Castenholz, R. W. (2003) The synthesis of the UV-screening pigment, scytonemin, and photosynthetic performance in isolates from closely related natural populations of cyanobacteria (*Calothrix* sp.). Environ Microbiol 5: 484–491. 10.1046/j.1462-2920.2003.00436.x

Frigaard, N.-U., Larsen, K. L., Cox, R. P. (1996) Spectrochromatography of photosynthetic pigments as a fingerprinting technique for microbial phototrophs. FEMS Microbiol Ecol 20: 69–77.

Gan F, and Bryant DA (2015a) Adaptive and acclimative responses of cyanobacteria to far-red light. Environ Microbiol 17: 3450–65.

Gan, F., Shen, G., Bryant, D. A. (2015b) Occurrence of far-red light photoacclimation (FaRLiP) in diverse cyanobacteria. Life 5: 4–24.

Gan F, Zhang S, Rockwell NC, Martin SS, Lagarias C, and Bryant DA (2014) Extensive remodeling of a cyanobacterial photosynthetic apparatus in far-red light. Science 345: 1312–1317.

Garcia-Pichel, F., Sherry, N. D., and Castenholz, R. W. (1992). Evidence for an ultraviolet sunscreen role of the extracellular pigment scytonemin in the terrestrial cyanobacterium *Chlorogloeopsis* sp. Photochem Photobiol 56: 17–23.

Geelen, B., Tack, N., and Lambrechts, A. (2013) A snapshot multispectral imager with integrated tiled filters and optical duplication. Advanced Fabrication Technologies for Micro/Nano Optics and Photonics VI, Proc SPIE 8613: 861314.

Gisriel, C. J. (2024) Recent structural discoveries of photosystems I and II acclimated to absorb far-red light. Biochim Biophys Acta Bioenerg 1865(3):149032. doi: 10.1016/j.bbabio.2024.149032.

Holst, G., and Grunwald, B. (2001) Luminescence lifetime imaging with transparent oxygen optodes. Sens Act B 74: 78–90.

Kirilovsky, D., and Kerfeld, C. A. (2016) Cyanobacterial photoprotection by the orange carotenoid protein. Nat Plants 2: 16180.

Klimant, I., Kühl, M., Glud, R. N. & Holst, G. (1997). Optical measurement of oxygen and temperature in microscale: strategies and biological applications. Sens Act B 38–39: 29–37.

Koren, K., and Kühl, M. (2018) Optical O2 sensing in aquatic systems and organisms. In: D. B. Papkovsky and R. Dmitriev (ed.), Quenched-phosphorescence detection of molecular oxygen: Applications in life sciences. Royal Society of Chemistry, Detection Science Series, 11: 145–174.

Krumbein, W. E. (1979) Photolithotropic and chemoorganotrophic activity of bacteria and algae as related to beachrock formation and degradation (Gulf of Aqaba, Sinai). Geomicrobiology Journal 1:139–203.

Kühl, M., Trampe, E., Mosshammer, M., Johnson, M., Larkum, A. W. D., and Koren, K. (2020) Substantial near-infrared radiation-driven photosynthesis of chlorophyll *f*-containing cyanobacteria in a natural habitat. eLife 9: e50871.

Li, Y., Scales, N., Blankenship, R. E., Willows, R. D., Chen, M. (2012) Extinction coefficient for red-shifted chlorophylls: Chlorophyll *d* and chlorophyll *f*. Biochim Biophys Acta 1817: 1292–1298.

Mascoli, V., Bersanini, L., and Croce, R. (2020) Far-red absorption and light-use efficiency trade-offs in chlorophyll *f* photosynthesis. Nat Plants 6, 1044–1053.

Mascoli, V., Bhatti, A.F., Bersanini, L. et al. (2022) The antenna of far-red absorbing cyanobacteria increases both absorption and quantum efficiency of Photosystem II. Nat Comm 13: 3562.

Mosshammer, M., Scholz, V. V., Holst, G., Kühl, M., and Koren, K. (2019) Luminescence lifetime imaging of O_2_ with a frequency domain based camera system. J Visual Exp 154: e60191.

Muramatsu, M., and Hihara, Y. (2012) Acclimation to high-light conditions in cyanobacteria: from gene expression to physiological responses. J Plant Res 125: 11–39.

Murray, B., Ertekin, E., Dailey, M., Soulier, N. T., Shen, G., Bryant, D. A., Perez-Fernandez, C., and DiRuggiero, J. (2022) Adaptation of cyanobacteria to the endolithic light spectrum in hyper-arid deserts. Microorganisms 10: 1198.

Niyogi, K. K., and Truong, T. B. (2013) Evolution of flexible non-photochemical quenching mechanisms that regulate light harvesting in oxygenic photosynthesis. Curr Opin Plant Biol 16: 307–314.

Santner, J., Larsen, M., Kreuzeder, A., Glud, R. N. (2015). Two decades of chemical imaging of solutes in sediments and soils - a review. Anal Chim Acta 878: 9–42.

Schindelin, J., Arganda-Carreras, I., Frise, E., Kaynig, V., Longair, M., Pietzsch, T., … Cardona, A. (2012). Fiji: an open-source platform for biological-image analysis. Nat Meth 9: 676–682. doi:10.1038/nmeth.2019

Schreiber U., Gademann, R., Bird, P., Ralph, P., Larkum, A.W.D., and Kühl, M. (2002) Apparent light requirement for activation of photosynthesis upon rehydration of dessicated beachrock microbial mats. J Phycology 38: 125–134.

Soulier, N., Laremore, T. N., and Bryant, D. A. (2020) Characterization of cyanobacterial allophycocyanins absorbing far-red light. Photosynth Res 145: 189–207.

Soulier, N., Walters, K., Laremore, T. N., Shen, G., Golbeck, J. H., and Bryant, D. A. (2022) Acclimation of the photosynthetic apparatus to low light in a thermophilic *Synechococcus* sp. strain. Photosynth Res 153: 21– 42.

Trampe, E., and Kühl, M. (2016) Chlorophyll *f* distribution and dynamics in cyanobacterial beachrock. J Phycol 52: 990–996.

Vousdoukas, M. I., Velegrakis, A.F., and Plomaritis T. A. (2007) Beachrock occurrence, characteristics, formation mechanisms and impacts. Earth-Science Rev 85: 23–46.

Zhao C, Gan F, Shen G, and Bryant DA (2015) RfpA, RfpB, and RfpC are the master control elements of far-red light photoacclimation (FaRLiP). Front Microbiol 6: 1–13.

Zieger, S., Mosshammer, M., Kühl, M., and Koren, K. (2020) Hyperspectral luminescence imaging in combination with signal deconvolution enables reliable multi-indicator based chemical sensing. ACS Sensors 6: 183–191.

